# *Arid1b* haploinsufficiency in cortical inhibitory interneurons causes cell-type-dependent changes in cellular and synaptic development

**DOI:** 10.1101/2024.06.07.597984

**Authors:** Alec H. Marshall, Danielle J. Boyle, Meretta A. Hanson, Devipriyanka Nagarajan, Noor Bibi, Alireza Safa, Aidan C. Johantges, Jason C. Wester

## Abstract

Autism spectrum disorder (ASD) presents with diverse cognitive and behavioral abnormalities beginning during early development. Although the neural circuit mechanisms remain unclear, recent work suggests pathology in cortical inhibitory interneurons (INs) plays a crucial role. However, we lack fundamental information regarding changes in the physiology of synapses to and from INs in ASD. Here, we used transgenic mice to conditionally knockout one copy of the high confidence ASD risk gene *Arid1b* from the progenitors of parvalbumin-expressing fast-spiking (PV-FS) INs and somatostatin-expressing non-fast-spiking (SST-NFS) INs. In brain slices, we performed paired whole-cell recordings between INs and excitatory projection neurons (PNs) to investigate changes in synaptic physiology. In neonates, we found reduced synaptic input to INs but not PNs, with a concomitant reduction in the frequency of spontaneous network events, which are driven by INs in immature circuits. In mature mice, we found a reduction in the number of PV-FS INs in cortical layers 2/3 and 5. However, changes in PV-FS IN synaptic physiology were cortical layer and PN cell-type dependent. In layer 5, synapses from PV-FS INs to subcortical-projecting PNs were weakened. In contrast, in layer 2/3, synapses to and from PV-FS INs and corticocortical-projecting PNs were strengthened, leading to enhanced feedforward inhibition of input from layer 4. Finally, we found a novel synaptic deficit among SST-NFS INs, in which excitatory synapses from layer 2/3 PNs failed to facilitate. Our data highlight that changes in unitary synaptic dynamics among INs in ASD depend on neuronal cell-type.

## INTRODUCTION

The genetic and environmental causes of autism spectrum disorder (ASD) are diverse but converge on a core set of behavioral and cognitive abnormalities. This suggests there may be shared pathophysiological mechanisms. Recent studies in monogenetic mouse models highlight that pathology in cortical inhibitory interneurons (INs) may be a strong contributor to ASD symptoms (Contractor et al., 2021; Nomura, 2021; Tang et al., 2021). Indeed, the same major IN classes are found across the cortex and engage in canonical circuit motifs (Tremblay et al., 2016; Tasic et al., 2018); thus, changes in the developmental trajectory of INs could lead to a constellation of diverse phenotypes, including social and sensory processing deficits. However, INs are diverse and the physiology of the synapses they send and receive depend on their identity and that of their partner. Thus, it is necessary to determine how these cell-type-specific synapses are altered, if changes are cell-autonomous, and if they cause ASD.

Most studies of IN pathology in ASD have focused on PV-FS cells, which include basket cells that target perisomatic compartments and chandelier cells that target the axon initial segment. PV+ IN numbers are reduced in cortical tissue from human patients diagnosed with ASD and in multiple mouse models, suggesting they may be particularly vulnerable (Godavarthi et al., 2014; Hashemi et al., 2017; Jung et al., 2017; Ariza et al., 2018; Kourdougli et al., 2023). This could be due to loss of PV expression in a subset of basket or chandelier cells (Filice et al., 2016). Indeed, PV expression is activity dependent (Patz et al., 2004) and PV+ INs are hypoactive in several ASD mouse models (Goel et al., 2018; Antoine et al., 2019). Alternatively, these cells may be lost, leading to fewer total INs in the cortex to regulate excitation (Jung et al., 2017; Kourdougli et al., 2023). Furthermore, in several ASD mouse models the frequency of miniature inhibitory postsynaptic currents (mIPSCs) recorded from excitatory projection neurons are reduced, suggesting deficits in PV-FS IN synapses (Jung et al., 2017; Antoine et al., 2019). Collectively, these changes may lead to an imbalance in the ratio of excitation to inhibition (E/I), which is hypothesized to contribute to ASD (Sohal and Rubenstein, 2019). However, few studies have investigated the physiology of unitary synaptic connections between PV-FS INs and specific postsynaptic cell-types. Thus, the extent to which changes in inhibitory synaptic strength generalize across cortical layers and regions or if there are changes in short-term synaptic dynamics that converge across ASD models are unknown. Furthermore, very little is known regarding the contributions of other IN subtypes, such as dendrite-targeting SST-NFS INs.

To investigate how pathology in INs may cause neurodevelopmental disorders, recent work has used a mouse model of *Arid1b* haploinsufficiency. ARID1B is a subunit of the Brg/Brahma-associated factor (BAF) chromatin remodeling complex (Wang et al., 1996; Wang et al., 2004) expressed in neurons beginning during embryonic development, and broadly associated with cell proliferation and differentiation (Moffat et al., 2019; Moffat et al., 2021a; Pagliaroli and Trizzino, 2021). In humans, mutations in *Arid1b* result in Coffin-Siris syndrome and nonsyndromic ASD, collectively referred to as ARID1B-related disorders (Santen and Clayton-Smith, 2014; van der Sluijs et al., 2019). Transgenic *Arid1b* haploinsufficient mice demonstrate several core ASD cognitive and behavioral deficits, including increased anxiety, impaired learning and memory, and abnormal social interactions, suggesting it is a robust model system to study neural circuit mechanisms (Celen et al., 2017; Jung et al., 2017; Ellegood et al., 2021; Moffat et al., 2021a; Kim et al., 2022). Furthermore, deficits in social communication are observed in neonatal mice (Celen et al., 2017; Kim et al., 2022), demonstrating *Arid1b* haploinsufficiency is also a powerful model of early neurodevelopmental disorders. Importantly, conditional knockout of one copy of *Arid1b* from neuronal progenitors of the medial ganglionic eminence (MGE), which produces both PV-FS and SST-NFS INs, is sufficient to cause behavioral phenotypes observed in germline *Arid1b* haploinsufficient mice (Jung et al., 2017). Furthermore, conditional *Arid1b* haploinsufficiency in either PV-FS or SST-NFS INs results in distinct sets of cognitive and behavioral deficits (Smith et al., 2020). However, how *Arid1b* haploinsufficiency alters their synaptic physiology to generate these abnormal behaviors remains unknown.

Here, we used transgenic mice to conditionally knockout one copy of *Arid1b* from interneuron progenitors of the MGE. We then used whole-cell recordings in brain slices to rigorously examine changes in synaptic physiology in both neonatal and adolescent mice among PV-FS INs, SST-NFS INs, and different PN subtypes. Our data reveal changes in synaptic strength and short-term plasticity that depend on developmental stage and neuronal cell-type.

## RESULTS

To investigate the impact of *Arid1b* haploinsufficiency on inhibitory interneurons (INs), we crossed Nkx2.1-Cre mice (Xu et al., 2008) to a transgenic line in which loxP sites flank exon 5 of the *Arid1b* allele (Celen et al., 2017). Nkx2.1 is a transcription factor expressed in progenitors of the medial ganglionic eminence (MGE), which give rise to fast spike parvalbumin-expressing cells (PV-FS INs) and non-fast spiking somatostatin-expressing cells (SST-NFS INs) (Anderson et al., 2001; Wichterle et al., 2001; Butt et al., 2005). This allowed us to conditionally knockout one copy of *Arid1b* from Nkx2.1- lineage INs during embryogenesis; we term these mice IN *Arid1b*(+/-). To identify these INs in tissue sections and to target them for recording, we further crossed these mice to a floxed tdTomato reporter line (Ai14). Previous studies using mice to investigate neuronal and synaptic physiology in ASD, including *Arid1b* haploinsufficiency, focused on the neocortex. Thus, to compare our data to those in the current literature, we performed most experiments in primary visual cortex. However, we assayed neonatal synapses in CA1 hippocampus due to the propensity of this region to generate spontaneous network activity in brain slices for high throughput screening of synaptic changes.

### *Arid1b* haploinsufficiency results in reduced number of PV-FS interneurons

Previously, Jung et al. (2017) found reduced numbers of GABA+ and PV+ cells in the neocortex of mice with germline, whole-body *Arid1b* haploinsufficiency. They observed the same phenotype in Nkx2.1-Cre;Arid1b(flox/+) mice, suggesting IN loss occurs by a cell-autonomous mechanism. However, they did not use a conditional reporter to identify mutant, Nkx2.1-Cre-expressing cells. This is important because the Nkx2.1-Cre mouse line does not express Cre recombinase in ∼30% of PV+ cells in layer 2/3 and 10% in layer 5 (Xu et al., 2008). Thus, to replicate and extend previous findings, we directly quantified the number of *Arid1b* haploinsufficient Nkx2.1-Cre-lineage cells identified by their expression of tdTomato and co-stained for PV (**Figure 1A**). Importantly, we did not find a reduction in the total numbers of Nkx2.1-Cre;tdTomato+ cells or PV+ cells (**Figure 1B**). However, we did find a reduction in the number of Nkx2.1-Cre;tdTomato+ cells that express PV (**Figure 1C**), which was due to loss of these cells in cortical layers 2/3 and 5 (**Figure 1D**). Thus, the total number of PV+ cells was not affected, despite the reduction in the percentage of Nkx2.1-Cre;tdTomato+ interneurons that express PV. Our data suggest that genetically normal PV+ cells (i.e., not represented in the Nkx2.1-Cre line) increase their number to compensate for the reduced number of *Arid1b* haploinsufficient Nkx2.1-Cre-lineage PV+ interneurons. Furthermore, because genetically normal PV+ interneurons were not reduced in number, our data suggest that loss of PV expression due to *Arid1b* haploinsufficiency is cell autonomous.

**Figure 1.**
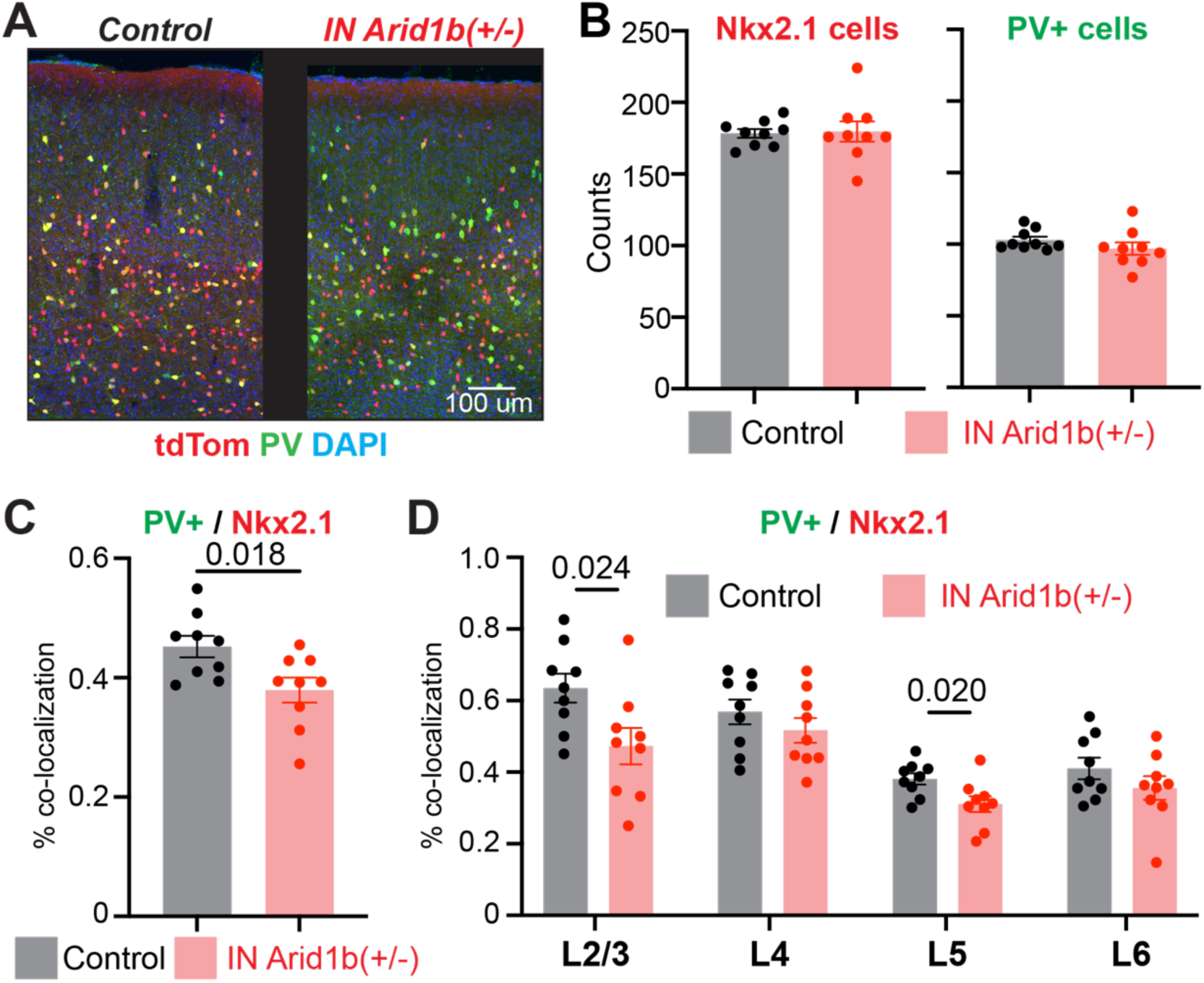
*Arid1b* haploinsufficiency results in fewer interneurons from the Nkx2.1-lineage that express PV. **A)** Representative example images of V1 from control and IN *Arid1b*(+/-) mice used to quantify tdTomato+ and PV+ cells. Quantifications are from n = 3 control and n = 3 mutant mice, 3 sections per mouse. **B)** No change in the total number of Nkx2.1-lineage interneurons (unpaired t test, t(16) = 0.1734, p = 0.86) or the total number of PV+ cells (Mann-Whitney U test, U = 21, p = 0.09). **C)** Fewer Nkx2.1-lineage interneurons express PV (unpaired t test, t(16) = 2.633). **D)** The reduction in Nkx2.1-lineage interneurons that express PV is limited to layers 2/3 (unpaired t test, t(16) = 2.484) and 5 (unpaired t test, t(16) = 2.574). For layer 4 (unpaired t test, t(16) = 1.048, p = 0.31); layer 6 (unpaired t test, t(16) = 1.237, p = 0.23).

Our data are broadly consistent with previous studies that found PV+ interneurons are reduced in *Arid1b* haploinsufficiency (Jung et al., 2017; Moffat et al., 2021b). However, because the total number of Nkx2.1-Cre-expressing cells were not changed in mutant mice (**Figure 1B**), it remained unclear if PV-FS INs remained but lacked PV expression, or if PV-FS INs were lost at the expense of SST-NFS INs. Indeed, *Arid1b* regulates ventral progenitor proliferation and survival, but it also can bind the *Pvalb* promotor to regulate transcription (Jung et al., 2017; Moffat et al., 2021b). To distinguish between these possibilities, we performed whole-cell patch clamp recordings of Nkx2.1-Cre;tdTomato+ INs in layer 2/3. In both control and IN *Arid1b*(+/-) mice, PV-FS and SST-NFS cells were easily distinguished based on their firing properties and input resistance (**Figure 2A**). Importantly, both interneuron classes were observed in IN *Arid1b*(+/-) mice and were not different from control. Specifically, in IN *Arid1b*(+/-) mice, PV-FS cells maintained a low input resistance (**Figure 2B**) and small voltage sag relative SST-NFS cells (**Figure 2C**). Furthermore, in response to depolarizing current steps at twice the threshold for action potentiation initiation, PV-FS cells attained higher firing rates than SST-NFS cells (**Figure 2D**) that did not attenuate in frequency (**Figure 2E**). Thus, *Arid1b* haploinsufficient PV-FS and SST-NFS interneurons observed in mature mice are physiologically normal; we did not find evidence of abnormal or “hybrid” cell-types. However, consistent with our immunohistochemistry data (**Figure 1**), we found a significant reduction in the ratio of FS to NFS cells sampled in IN *Arid1b*(+/-) mice relative to controls (**Figure 2F**). Thus, our data suggest that *Arid1b* haploinsufficiency causes a loss of PV-FS interneurons, rather than down-regulation of PV.

**Figure 2.**
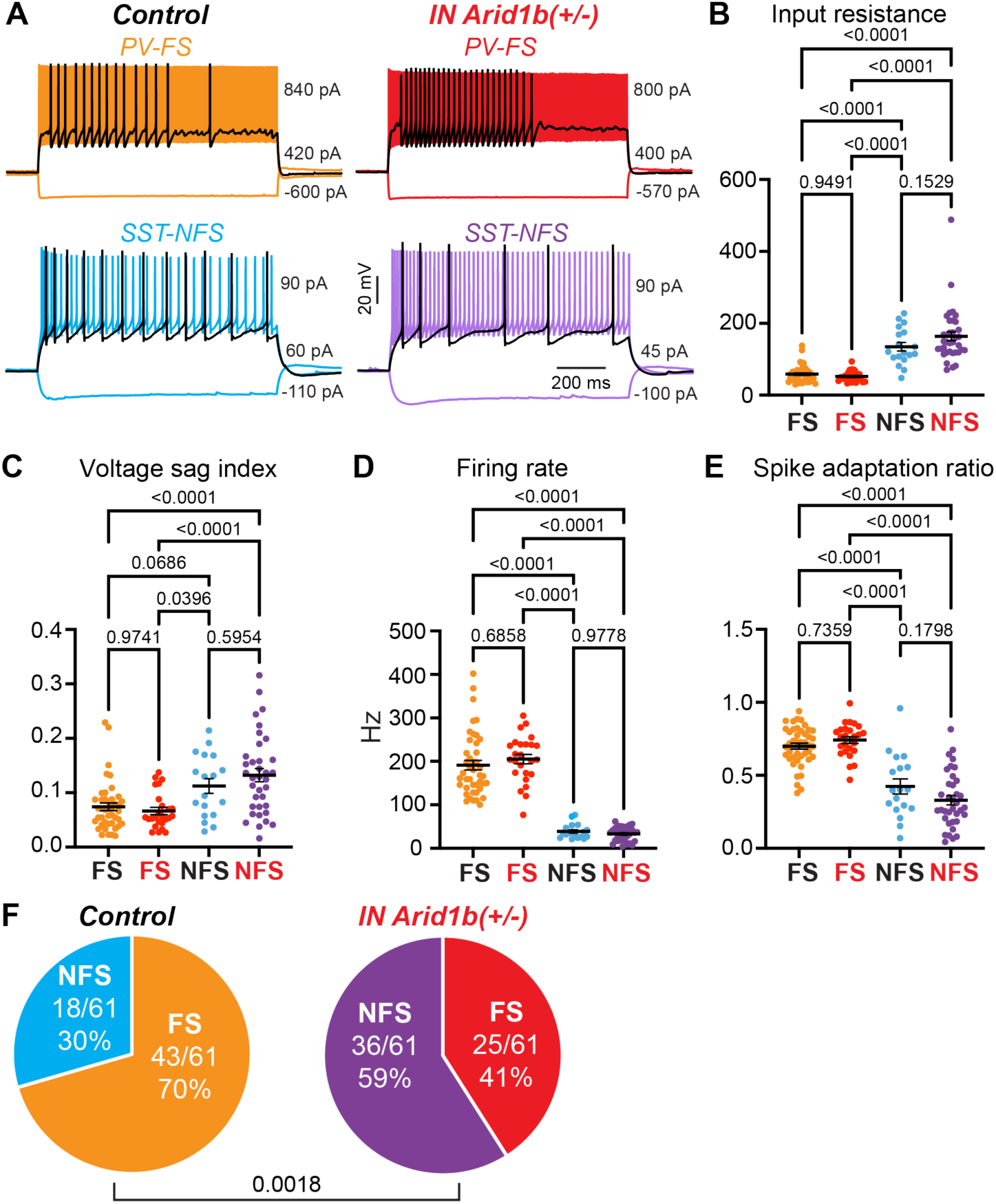
*Arid1b* haploinsufficiency results in fewer FS interneurons among Nkx2.1-lineage cells. **A)** Representative example recordings of PV-FS and SST-NFS interneurons from control and IN *Arid1b*(+/-) mice. All cells were biased to a resting membrane potential of -70 mV. Max depolarizing current is at twice the threshold for action potential initiation. Middle depolarizing current (black) is just above threshold to highlight firing properties. Hyperpolarizing current pulse is to steady-state membrane voltage of -90 mV to measure voltage sag. **B)** Input resistance. Two-way ANOVA, main effect of cell-type (F[1, 118] = 106.5, p < 0.0001), no main effect of genotype (F[1, 118] = 1.546, p = 0.2162). **C)** Voltage sag. Two-way ANOVA, main effect of cell-type (F[1, 118] = 24.38, p < 0.0001), no main effect of genotype: (F[1, 118] = 0.3411, p = 0.5603). **D)** Firing rate at twice current threshold for action potentials. Two-way ANOVA, main effect of cell-type (F[1, 118] = 277.6, p < 0.0001), no main effect of genotype: (F[1, 118] = 0.1794, p = 0.6727). **E)** Firing adaptation ratio at twice current threshold for action potentials. Two-way ANOVA, main effect of cell-type (F[1, 118] = 121.6, p < 0.0001), no main effect of genotype (F[1, 118] = 0.7476, p = 0.3890). **F)** Ratios of FS to NFS INs observed in control vs. IN *Arid1b*(+/-) mice. Fisher’s exact test. For (B) – (E), Control FS n = 43, Control NFS n = 18, *Arid1b*(+/-) FS n = 25, *Arid1b*(+/-) FS n = 36, post hoc analysis is Tukey’s multiple comparisons test.

### Synaptic input to INs is weakened and the frequency of early spontaneous network events are reduced in neonates

In addition to neuronal progenitor proliferation, *Arid1b* is necessary for synapse development in both INs and PNs (Ka et al., 2016; Celen et al., 2017; Jung et al., 2017; Kim et al., 2022). Neonatal mice with *Arid1b* haploinsufficiency demonstrate altered vocal communication (Celen et al., 2017; Kim et al., 2022), suggesting disruptions in synaptic connectivity and physiology begin during synaptogenesis. In the rodent neocortex and hippocampus, synapses among INs and PNs form during the end of the first postnatal week (Pangratz-Fuehrer and Hestrin, 2011; Wester and McBain, 2016). This coincides with the occurrence of spontaneously generated network activity thought to be necessary for cellular and circuit maturation (Ben-Ari, 2001; Allene et al., 2008; Blankenship and Feller, 2010). In brain slices, these spontaneous events are called giant depolarizing potentials (GDPs) (Ben-Ari et al., 1989). GDPs travel as a wave, engage the entire local network of neurons for several hundred milliseconds, and occur robustly in hippocampal region CA1 (Wester and McBain, 2016). At this developmental stage GABA is depolarizing (Ben-Ari et al., 2007), and we previously showed that MGE-derived INs play a crucial role in generating GDPs (Wester and McBain, 2016). Thus, GDPs are a powerful tool to assay synaptic connectivity and strength in immature circuits.

In slices of neonatal CA1 hippocampus, we targeted a Nkx2.1-Cre;tdTomato+ interneuron and neighboring pyramidal cell for dual whole-cell patch clamp recording (**Figure 3A**). We used internal recording solution with a chloride reversal potential of -27 mV and voltage-clamped neurons at -70 mV (**Figure 3A**). This allowed us to record the total synaptic current during GDPs to assay changes in synaptic connectivity and strength in immature INs with *Arid1b* haploinsufficiency and neighboring pyramidal cells. In control mice, spontaneous GDPs occurred synchronously in INs and pyramidal cells with a frequency of ∼0.08 Hz, as observed previously (Wester and McBain, 2016) (**Figure 3B, left**). In IN *Arid1b*(+/-) mice, GDPs remained synchronous between INs and pyramidal cells (**Figure 3B, right**), but occurred with a significantly lower frequency relative to controls (**Figures 3B and 3C**). During individual GDPs, inward currents observed in both INs and pyramidal cells reached comparable peak amplitudes of several hundred picoamps (**Figure 3D, left**). However, in IN *Arid1b*(+/-) mice we noticed that peak synaptic currents were weaker in INs relative to simultaneously recorded neighboring pyramidal cells (**Figure 3D, right**). To quantify changes in synaptic currents, we measured the charge transfer (integral of the current) during individual GDPs. In IN *Arid1b*(+/-) mice, we found that GDP charge transfer was significantly lower in INs, but was not altered in pyramidal cells (**Figure 3E**). Changes in charge transfer could occur due to differences in synaptic current amplitude, GDP duration, or both. We found that maximum current amplitude was significantly reduced in INs, but not pyramidal cells (**Figure 3F**). However, the duration of GDPs was not different in either INs or pyramidal cells in both conditions (**Figure 3G**). Thus, in IN *Arid1b*(+/-) mice, recordings from the perspective of pyramidal cells indicate that when GDPs are generated within the network, they proceed normally: they are not truncated in duration and the strength of synaptic inputs are not altered. However, *Arid1b* haploinsufficiency causes cell-autonomous weakening of synaptic inputs to INs during early postnatal development, which can be observed during GDPs (**Figure 3H**).

**Figure 3.**
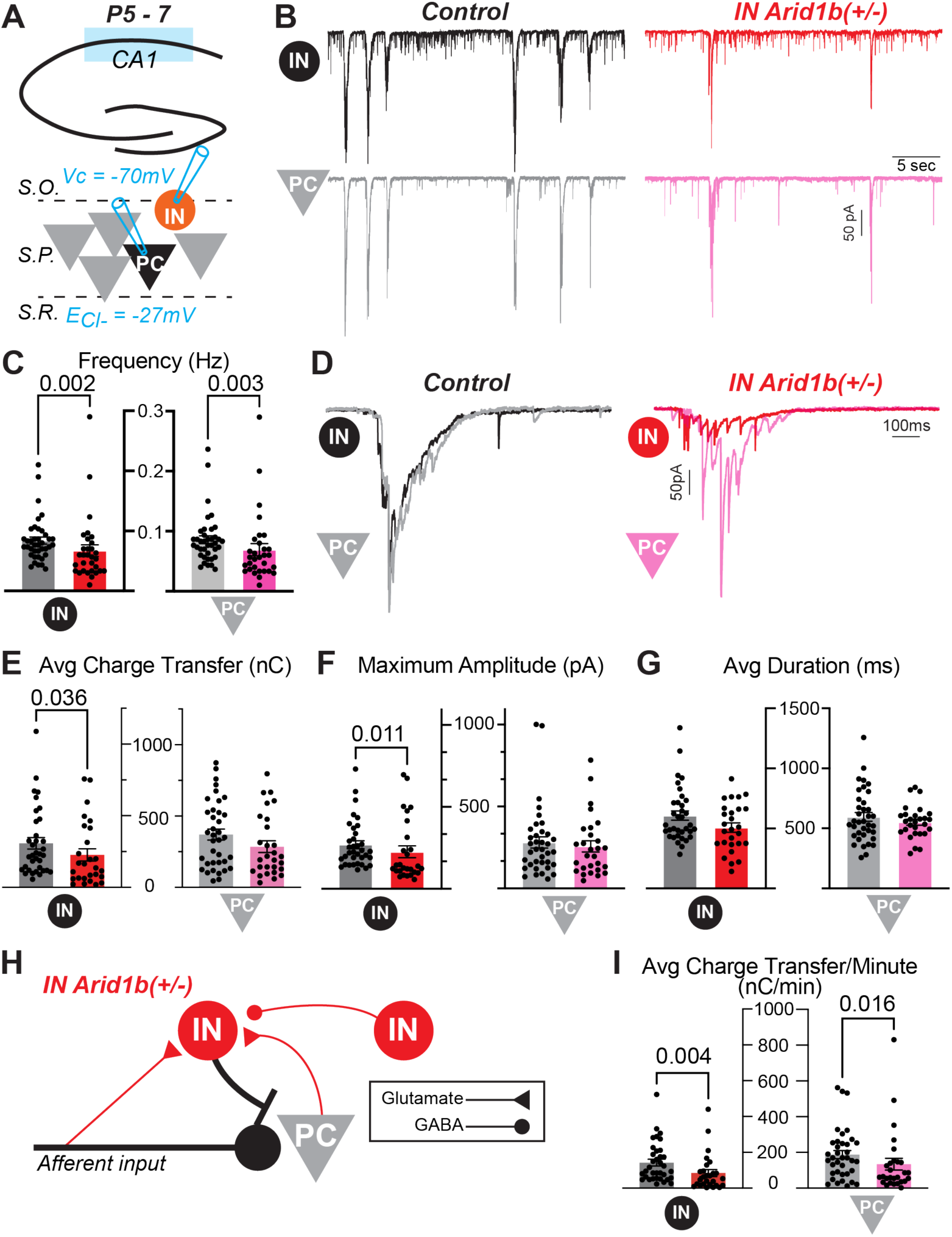
*Arid1b* haploinsufficiency in Nkx2.1-lineage interneurons results in weaker input selectively to interneurons and reduced frequency of GDPs during the end of the first postnatal week. **A)** Experimental configuration. In slices of neonatal CA1 hippocampus, interneurons and pyramidal cells were patched in pairs. Both were voltage-clamped at -70 mV using an internal recording solution with a chloride reversal potential of -27 mV. Thus, all synaptic currents during GDPs are inward. **B)** Representative recordings of spontaneous GDPs. Note the lower frequency in IN Arid1b(+/-) mice. **C)** Quantification of GDP frequencies observed in interneurons and pyramidal cells. IN control, n = 39; IN mutant, n = 31; PC control, n = 39; PC mutant, n = 31. Mann-Whitney tests. **D)** Representative recordings of individual GDPs. Note in control the amplitude and duration of GDPs recorded in interneurons and pyramidal cells are equivalent, but in IN Arid1b(+/-) mice the amplitude of GDPs recorded in interneurons is reduced. **E)** Quantification of the average charge transfer of GDPs recorded in individual interneurons and pyramidal cells. IN control, n = 35; IN mutant, n = 27; PC control, n = 37; PC mutant, n = 28. Mann-Whitney tests. For pyramidal cells, p = 0.123. **F)** Quantification of the average maximum amplitude of GDPs recorded in individual interneurons and pyramidal cells. IN control, n = 35; IN mutant, n = 27; PC control, n = 37; PC mutant, n = 28. Mann-Whitney tests. For pyramidal cells, p = 0.626. **G)** Quantification of the average duration of GDPs recorded in individual interneurons and pyramidal cells. IN control, n = 35; IN mutant, n = 27; PC control, n = 37; PC mutant, n = 28. For interneurons, Mann-Whitney test, p = 0.133. For pyramidal cells, unpaired t test, p = 0.348. **H)** Schematic illustrating that in IN Arid1b(+/-) there is a reduction in synaptic input to interneurons but not pyramidal cells. **I)** Quantification of the average charge transfer per minute from GDPs recorded in individual interneurons and pyramidal cells. IN control, n = 35; IN mutant, n = 27; PC control, n = 37; PC mutant, n = 28. Mann-Whitney tests.

Finally, MGE-derived INs are critical contributors to GDP initiation (Wester and McBain, 2016), thus, weak synaptic drive to these cells is likely the cause of reduced GDP frequency in IN *Arid1b*(+/-) mice (**Figure 3C**). In vivo, depolarization during spontaneous network activity induces calcium influx that contributes to neuronal and synaptic development and interneuron survival (Ben-Ari, 2001; Spitzer, 2006; Denaxa et al., 2018; Priya et al., 2018; Wong et al., 2018). We found that lower GDP frequency in IN *Arid1b*(+/-) mice leads to a reduction in the average charge transfer per minute in both INs and pyramidal cells (**Figure 3I**). Thus, *Arid1b* haploinsufficiency in INs may have long-term consequences for neuronal and circuit development.

### Inhibitory synapses from PV-FS INs to pyramidal tract-type PNs in layer 5 are weakened in mature mice

Previous studies using miniature inhibitory postsynaptic currents (mIPSCs) to assay changes in synaptic connectivity and physiology in germline, whole-body *Arid1b* haploinsufficient mice produced conflicting results. Jung et al. (2017) found reduced frequency of mIPSCs recorded in layer 5 PNs, but Kim et al. (2022) found no difference in either layers 2/3 or 5. In layer 5, PNs are a mix of intratelencephalic (IT)-type and pyramidal-tract (PT)-type cells (Harris and Shepherd, 2015) that form unique synaptic connections dependent on their cell-type. In paired whole-cell recordings, PV-FS INs form synaptic connections with PT-type PNs at a higher rate than IT-type PNs (Lee et al., 2014). Thus, to assay PV-FS synaptic connectivity and physiology in layer 5, we injected a retrograde tracer into the superior colliculus (**Figure 4A**) to label PT-type PNs in V1 (**Figure 4B**). We then performed paired whole-cell recordings in layer 5 between PT-type PNs and Nkx2.1-Cre;tdTomato+ cells identified as FS (**Figure 4C**). We found that the probability of finding synaptic connections from PV-FS INs to PT-type PNs or from PT-type PNs to PV-FS INs were equivalent in control and IN *Arid1b*(+/-) mice (**Figure 4D**). However, we found that the average inhibitory synaptic currents observed in connected pairs were smaller in *Arid1b*(+/-) mice (**Figure 4E**). To quantify differences in synaptic physiology, we analyzed synaptic potency (the average IPSC amplitude for trials in which a presynaptic action potential evoked vesicle release) and synaptic failure rate (the percentage of trials in which a presynaptic action potential did not evoke vesicle release) (Stevens and Wang, 1995). We found that the potency of IPSCs recorded in PT-type PNs were significantly reduced in *Arid1b*(+/-) mice (**Figure 4F**). However, synaptic failure rate was not different between conditions (**Figure 4G**). Thus, our data are consistent with previous work showing that inhibition is weaker in layer 5 of *Arid1b* haploinsufficient mice. Furthermore, our data suggest this is due to cell autonomous mechanisms in PV-FS INs.

**Figure 4.**
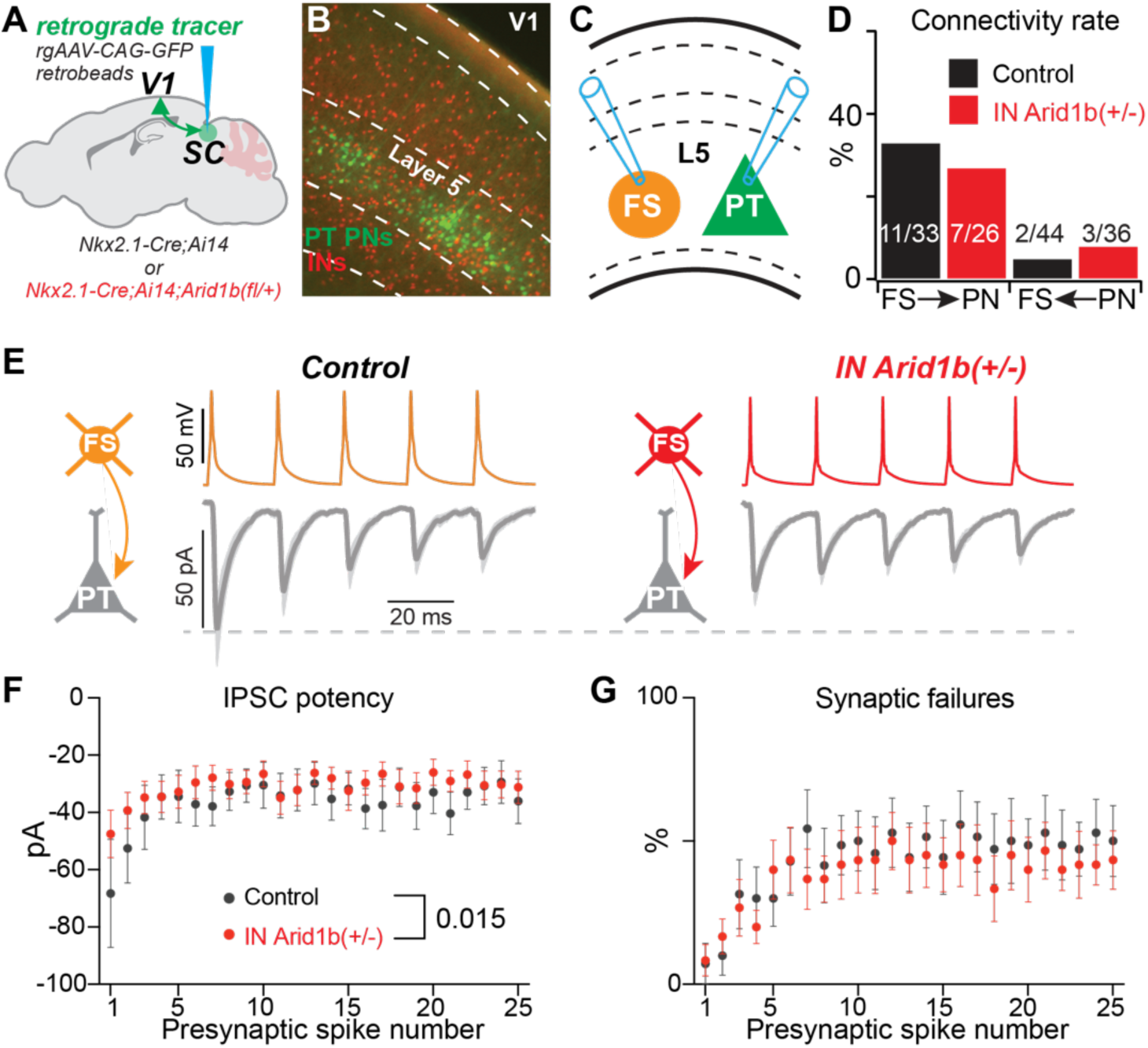
*Arid1b* haploinsufficiency in Nkx2.1-lineage interneurons results in weaker inhibitory synapses from FS interneurons to layer 5 PT-type PNs. **A)** Schematic of retrograde tracer injection site in the superior colliculus (SC) to label layer 5 pyramidal-tract type PNs in V1. **B)** Representative image of retrogradely labeled PT-type PNs and Nkx2.1-lineage interneurons labeled with tdTomato. **C)** Configuration for paired whole-cell recordings to test for unitary synaptic connections. **D)** Synaptic connectivity rates are comparable in control and IN *Arid1b*(+/-) mice. For FS IN to PN, Fisher’s exact test (p = 0.777). For PN to FS IN, Fisher’s exact test (p = 0.653). **E)** Average inhibitory postsynaptic currents recorded in control and IN *Arid1b*(+/-) mice. Control, n = 11 pairs; IN *Arid1b*(+/-), n = 7 pairs. **F)** Quantification of IPSC potency from observed synaptic connections. N’s same as (D). Main effect of genotype, two-way ANOVA: F(1,270) = 5.992. **G)** Percentage of trials in which a presynaptic action potential in a FS IN failed to evoke vesicle release at synapse onto a PT-type PC. Main effect of genotype, two-way ANOVA: F(1,275) = 2.843, p = 0.093.

### Inhibitory synapses between PV-FS INs and intratelencephalic-type PNs in layer 2/3 are strengthened

Next, we investigated synaptic connections between PV-FS INs and PNs in layer 2/3, which are all IT-type (**Figure 5A**). Like in layer 5, the probability of observing synaptic connections from PV-FS INs to PNs were equivalent in control and IN *Arid1b*(+/-) mice (**Figure 5B**). However, we noted a small increase in the average amplitude of IPSCs in IN *Arid1b*(+/-) mice (**Figure 5C**). Indeed, we found that the potency of inhibitory synapses was significantly greater throughout the PV-FS presynaptic spike train (**Figure 5D**), although synaptic failure rates were normal (**Figure 5E**). Thus, in contrast to layer 5, inhibition from PV-FS cells in layer 2/3 is enhanced. Next, we investigated synaptic connections from PNs to PV-FS INs. Like those observed for inhibitory synapses, the synaptic connectivity rates were not different between conditions (**Figure 5F**). However, the average amplitudes of excitatory currents were larger in IN *Arid1b*(+/-) mice (**Figure 5G**). This was due to significantly greater EPSC potency (**Figure 5H**); excitatory synaptic failure rates were equivalent (**Figure 5I**). To determine if these differences in IPSC and EPSC potency were functionally relevant, we investigated feedforward inhibition of layer 4 excitatory input to layer 2/3 PNs. For these experiments, we performed a stimulation protocol previously used to investigate feedforward inhibition of other mouse models of ASD (Antoine et al., 2019). We placed a stimulating electrode in layer 4, and voltage-clamped a PN at -70 mV to measure evoked EPSCs and then at 0 mV to measure evoked feedforward IPSCs (**Figure 6A**). For each recorded PN, we first found the smallest stimulation intensity necessary to evoke an EPSC (termed Eε), and then generated input-output curves for excitation and inhibition at 1 – 1.5 times this intensity (**Figure 6B**). We found that the strength of excitatory synaptic input from layer 4 to layer 2/3 PNs was equivalent in control and IN *Arid1b*(+/-) mice (**Figure 6C**). However, consistent with our findings from paired whole-cell recordings (**Figure 5**), we found that feedforward inhibition of layer 4 input to layer 2/3 PNs was greater in IN *Arid1b*(+/-) mice (**Figure 6D**). Thus, our data show that, as in in layer 5, changes in inhibitory synaptic strength occur cell-autonomously. However, these changes are not uniform across the cortical circuits.

**Figure 5.**
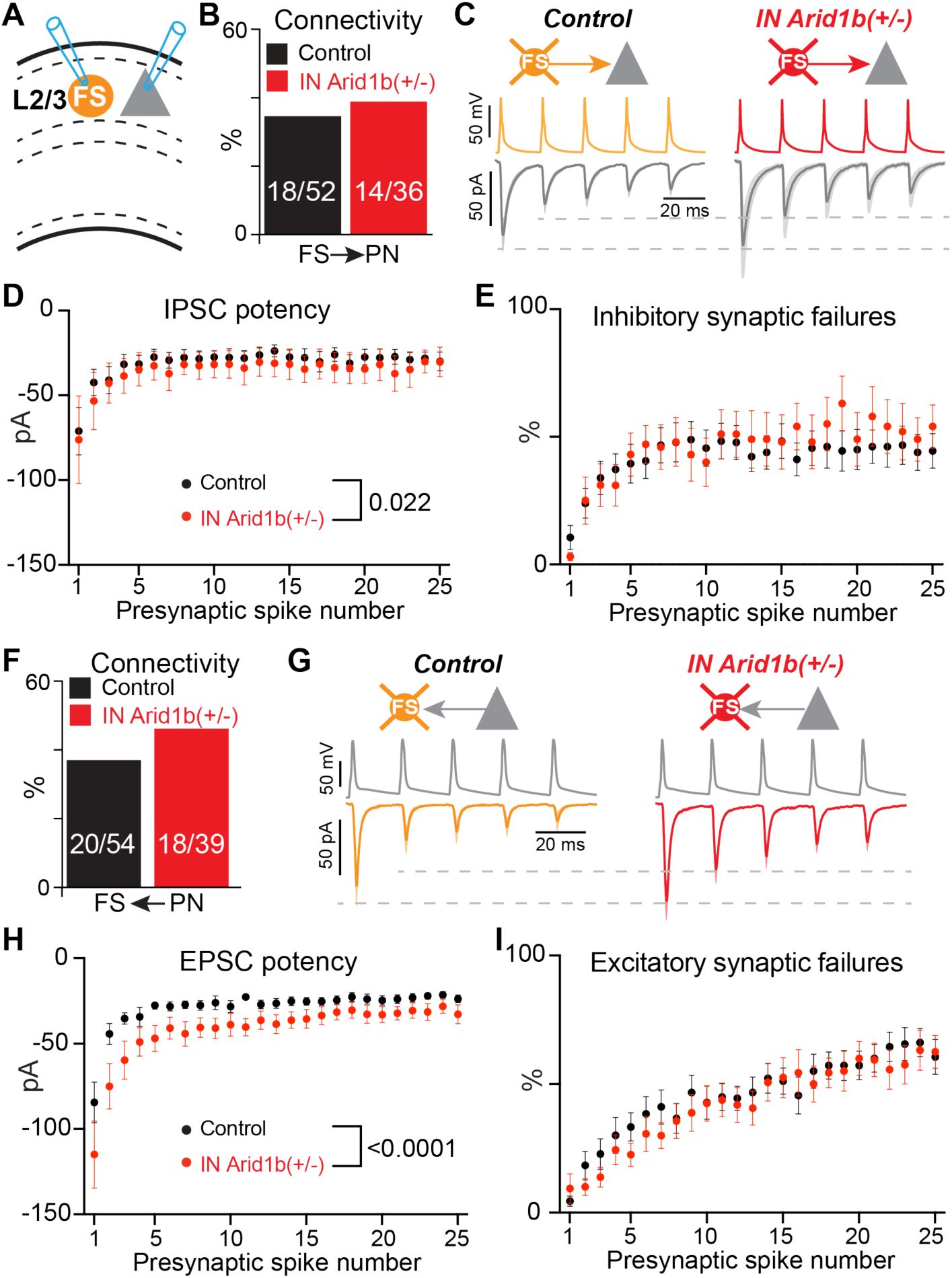
*Arid1b* haploinsufficiency in Nkx2.1-lineage interneurons results in stronger synaptic from FS interneurons to layer 2/3 PNs. **A)** Configuration for paired whole-cell recordings to test for unitary synaptic connections. **B)** Inhibitory synaptic connectivity rates are comparable in control and IN *Arid1b*(+/-) mice; Fisher’s exact test (p = 0.822). **C)** Average inhibitory postsynaptic currents recorded in control and IN *Arid1b*(+/-) mice. Control, n = 18 pairs; IN *Arid1b*(+/-), n = 10 pairs. **D)** Quantification of IPSC potency from observed synaptic connections. N’s same as (C). Main effect of genotype, two-way ANOVA: F(1,627) = 5.251. **E)** Percentage of trials in which a presynaptic action potential in a FS IN failed to evoke vesicle release at synapse onto a layer 2/3 PC. Main effect of genotype, two-way ANOVA: F(1,650) = 2.189, p = 0.14. **F)** Excitatory synaptic connectivity rates are comparable in control and IN *Arid1b*(+/-) mice; Fisher’s exact test (p = 0.400). **G)** Average excitatory postsynaptic currents recorded in control and IN *Arid1b*(+/-) mice. Control, n = 18 pairs; IN *Arid1b*(+/-), n = 16 pairs. **H)** Quantification of EPSC potency from observed synaptic connections. N’s same as (G). Main effect of genotype, two-way ANOVA: F(1,777) = 64.21. **I)** Percentage of trials in which a presynaptic action potential in a PN failed to evoke vesicle release at synapse onto a FS IN. Main effect of genotype, two-way ANOVA: F(1,800) = 3.626, p = 0.0573.

**Figure 6.**
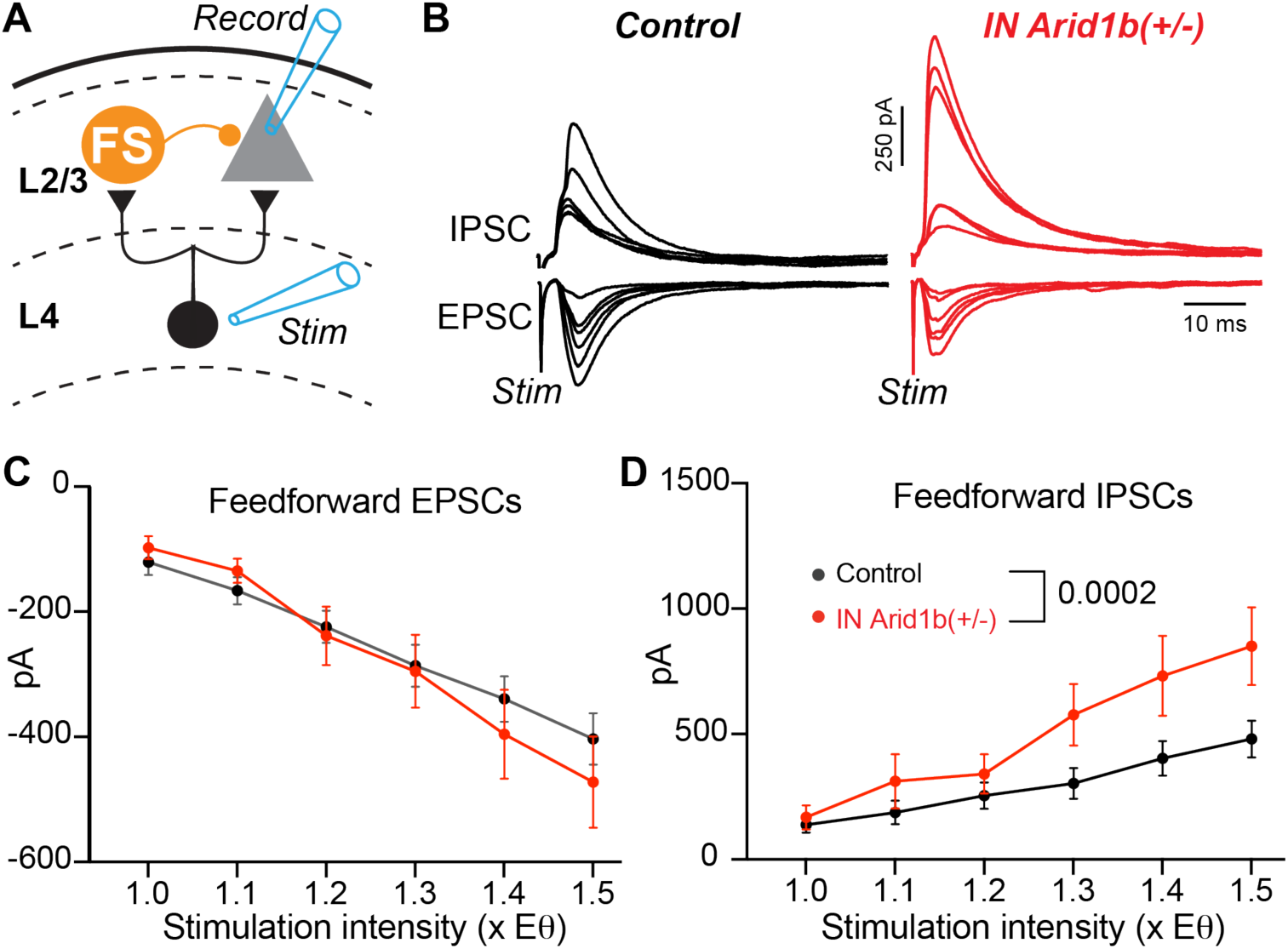
*Arid1b* haploinsufficiency in Nkx2.1-lineage interneurons results in increased feedforward inhibition of layer 4 input to layer 2/3 PNs. **A)** Recording and stimulation configuration. **B)** Representative examples of evoked EPSCs and IPSCs recorded in a layer 2/3 PN in a control and IN *Arid1b*(+/-) mouse. **C)** Input-output curve for evoked EPSC amplitudes. The strength of evoked EPSCs were not different between control (n = 20 cells) and IN Arid1b(+/-) (n = 20 cells) conditions. Main effect of genotype, two-way ANOVA: F(1,228) = 0.3918, p = 0.532. **D)** Input-output curve for evoked IPSC amplitudes. The strength of evoked IPSCs were significantly greater in IN *Arid1b*(+/-) mice (n = 20 cells) than in control mice (n = 20 cells). Main effect of genotype, two-way ANOVA: F(1,228) = 14.15.

### Excitatory synapses from PNs to SST-NFS INs have impaired facilitation

Finally, no studies have investigated if *Arid1b* haploinsufficiency impacts the physiology of synapses between PNs and other IN subtypes. Thus, we used paired whole-cell recordings to investigate synapses between PNs and Nkx2.1-Cre;tdTomato+ cells identified as NFS in layer 2/3 (**Figure 7A**). Although we did not observe inhibitory synaptic connections, we found excitatory connections from PNs to SST-NFS INs with high probability in both control and IN *Arid1b*(+/-) mice (**Figure 7B**). Excitatory synapses to SST-NFS INs in both the neocortex and hippocampus demonstrate striking facilitation during trains of presynaptic action potentials (Reyes et al., 1998; Gibson et al., 1999; Losonczy et al., 2002; Sun et al., 2009; Sylwestrak and Ghosh, 2012; Urban-Ciecko et al., 2018; Stachniak et al., 2019). In control mice, we consistently observed this form of short-term synaptic plasticity in paired recordings (**Figure 7C, left**). However, in IN *Arid1b*(+/-) mice we found that synaptic facilitation was severely impaired (**Figure 7C, right**). In control mice, synaptic potency increased as a function of presynaptic spike number (**Figure 7D, blue**), with a concomitant decrease in synaptic failure rate (**Figure 7E, blue**). In contrast, in IN *Arid1b*(+/-) mice, synaptic potency was weak and remained static throughout the presynaptic spike train (**Figure 7D, purple**). However, we observed a decrease in synaptic failure rate throughout the train that was comparable to control (**Figure 7E, purple**). As described in the Discussion, the dissociation of the impact of *Arid1b* haploinsufficiency on potency and failure rate may be due differential changes in the activation of presynaptic glutamate receptors. This novel synaptic deficit in SST-NFS INs may impair feedback and lateral inhibition in the neocortex in ARID1B-related disorders.

**Figure 7.**
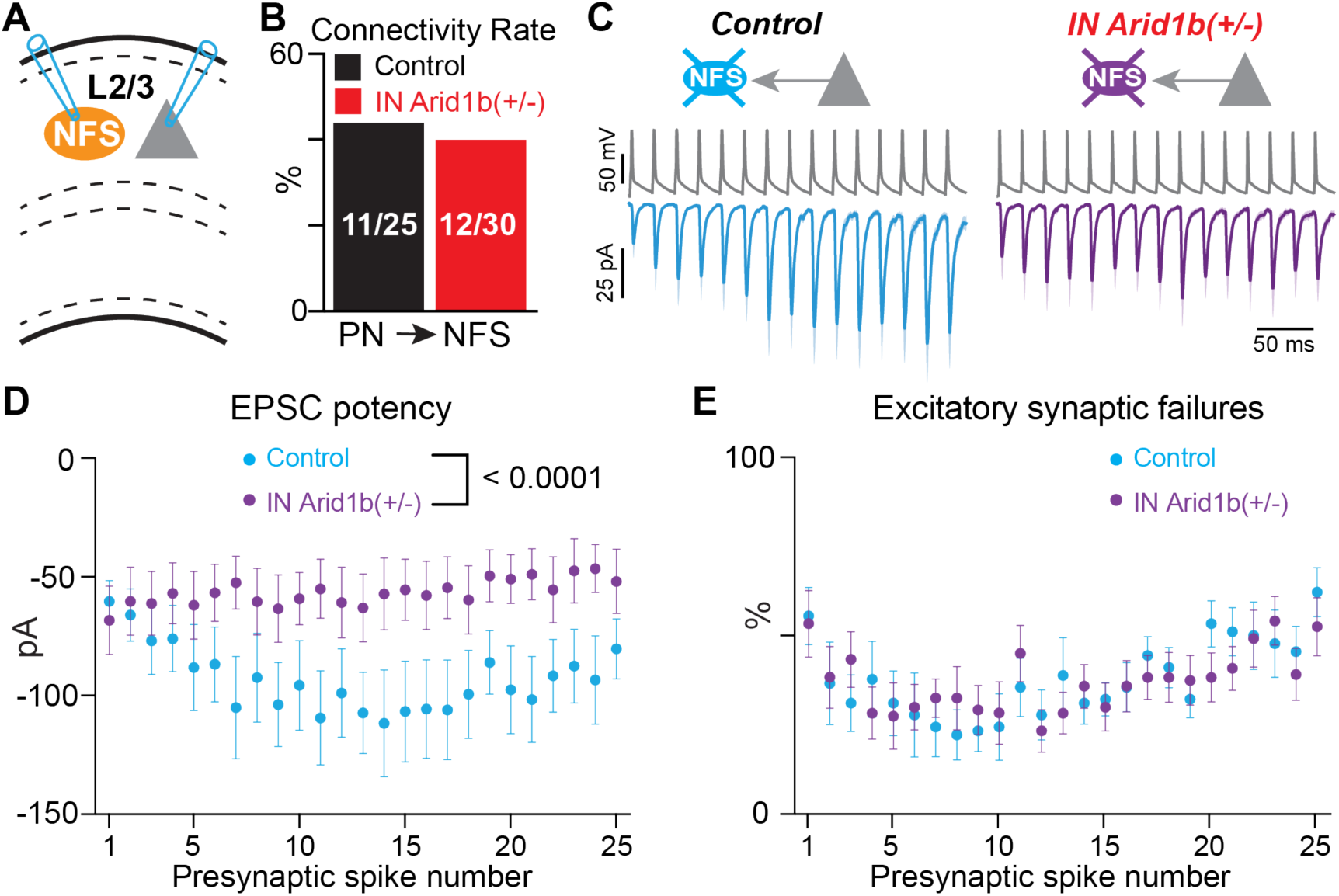
*Arid1b* haploinsufficiency in Nkx2.1-lineage interneurons results in weaker excitatory synapses from layer 2/3 PNs to NFS INs. **A)** Configuration for paired whole-cell recordings to test for unitary synaptic connections. **B)** Excitatory synaptic connectivity rates are comparable in control and IN *Arid1b*(+/-) mice; Fisher’s exact test (p = 0.790). **C)** Average excitatory postsynaptic currents recorded in control and IN *Arid1b*(+/-) mice. Control, n = 9 pairs; IN *Arid1b*(+/-), n = 12 pairs. **D)** Quantification of EPSC potency from observed synaptic connections. N’s same as (C). Main effect of genotype, two-way ANOVA: F(1,469) = 73.77. **E)** Percentage of trials in which a presynaptic action potential in a PN failed to evoke vesicle release at synapse onto a NFS IN. Main effect of genotype, two-way ANOVA: F(1,475) = 0.2084, p = 0.648.

## DISCUSSION

### Causes and consequences of reduced PV-FS interneuron density

In sections of prefrontal cortex collected from postmortem tissue in individuals diagnosed with autism, Hashemi et al. (2017) found reduced density of PV+ cells relative to those that express calbindin or calretinin. These data were the first to suggest that PV+ INs are selectively vulnerability in humans with ASD. Later work demonstrated similar loss of PV+ INs in cortical sections taken from patients diagnosed with Fragile X Syndrome (FXS) (Kourdougli et al., 2023). Reduction of PV+ INs is also commonly found in multiple mouse models of ASD (Contractor et al., 2021), including *Arid1b* haploinsufficiency (Jung et al., 2017). Importantly, reduced immunostaining for PV could be due to loss of PV expression in existing cells, loss of cells characterized by PV expression, or both (Filice et al., 2020). *Arid1b* haploinsufficiency is an interesting model to investigate these possibilities because there is evidence for reduced proliferation and increased apoptosis of IN progenitors, which suggests reduced cell number; however, *Pvalb* transcription is also reduced, suggesting the cells remain but lack PV (Jung et al., 2017; Moffat et al., 2021b). We used the Nkx2.1-Cre mouse line to conditionally knockout one copy of *Arid1b* from the progenitors of PV-FS and SST-NFS INs. Furthermore, we tagged Nkx2.1-Cre-lineage cells with tdTomato to allow us to quantify their numbers in the cortex and to target them for whole-cell recording regardless of their PV expression level. We found a reduction in the number of Nkx2.1-Cre;tdTomato+ cells that express PV and fewer INs with FS electrophysiological properties relative to those defined as NFS. Thus, although we cannot rule out that PV expression was down-regulated in a subset of INs, our data suggest that PV-FS INs are indeed lost in *Arid1b* haploinsufficiency.

Future PV-FS and SST-NFS INs can arise from the same a progenitor, thus it is somewhat surprising that only PV-FS cells are lost (Brown et al., 2011; Ciceri et al., 2013; Harwell et al., 2015; Mayer et al., 2015; Bandler et al., 2016). One possibility is that *Arid1b* haploinsufficiency differentially impacts apical versus basal progenitors. SST-NFS INs are born earlier and arise from apical progenitors, while PV-FS INs are born later and primarily produced by basal progenitors (Petros et al., 2015). Thus, it has been hypothesized that apical progenitors can undergo asymmetrical division to produce an SST-NFS IN and a basal progenitor that will give rise to a PV-FS IN (Bandler et al., 2016). Because PV-FS INs, but not SST-NFS INs, are lost in *Arid1b* haploinsufficient mice, our data suggest that basal progenitors during later phases of embryogenesis are selectively vulnerable to reduced *Arid1b* expression.

During early postnatal development, we found that *Arid1b* haploinsufficiency leads to reduced synaptic input to Nkx2.1-lineage INs. This may be a complementary or alternative mechanism by which PV-FS INs are lost. INs are initially generated in excessive numbers, and then undergo activity-dependent programmed cell death beginning at postnatal day 5 (Southwell et al., 2012; Denaxa et al., 2018; Wong et al., 2018; Duan et al., 2020). Thus, weak synaptic input to *Arid1b* haploinsufficient INs, coupled with reduced frequency of spontaneously generated network activity, is expected to impact their survival. Our data suggest immature, future PV-FS INs are particularly vulnerable; their numbers were reduced at the expense of increased numbers of SST-NFS INs. Such a redistribution of IN subtypes during early postnatal development has been observed previously. For example, reducing the number of medial ganglionic eminence-derived PV+ and SST+ INs results in a compensatory increase in the number of INs derived from the caudal ganglionic eminence and vice versa (Denaxa et al., 2018). Our data suggest a similar mechanism occurs among PV+ and SST+ INs. Finally, our data resemble those recently described by Kourdougli et al. (2023) in a FXS mouse model. They used the Nkx2.1-Cre line to conditionally knockout *Fmr1* and observed reduced activity of Nkx2.1-lineage interneurons *in vivo* at postnatal day 6, with a concomitant reduction in the number of PV+ INs in mature mice. Furthermore, like our study, they found an increase in the density of SST INs.

Importantly, although multiple studies have explored the mechanisms by which PV-FS INs are lost in ASD, the functional consequences are unclear. That is, is reduced density of PV-FS INs sufficient to generate ASD cognitive and behavioral phenotypes? Kourdougli et al. (2023) found that they could rescue the number of PV+ INs in somatosensory cortex by chemogenetically depolarizing Nkx2.1-lineage INs between postnatal days 5 – 10. However, this was insufficient to rescue abnormal responses of neurons to whisker stimuli observed in FXS mice. Thus, loss of PV+ INs may be a symptom rather than cause of ASD; changes in synaptic connectivity and physiology may play a more important role.

### Cell-type-specific changes in synapses to and from PV-FS interneuron and the balance of excitation to inhibition

A reduction in the number of PV-FS INs is expected to result in reduced inhibitory input to PNs. However, previous studies produced conflicting results regarding changes in synaptic connectivity and physiology in *Arid1b* haploinsufficient mice. In layer 5, Jung et al. (2017) found fewer inhibitory synaptic puncta and reduced frequency of mIPSCs recorded from PNs. In contrast, Kim et al. (2022) found no change in mIPSC frequency or amplitude in layers 2/3 or 5. We used paired whole-cell recordings between specific pre- and postsynaptic neurons to assay synaptic connectivity and physiology of evoked transmitter release. In layer 5, we found no difference in connectivity rate between PV-FS INs and PT-type PNs, however, we found a small but significant decrease in the potency of unitary inhibitory synapses. Thus, our data are consistent with Jung et al. (2017) that inhibition is weakened in infragranular cortical layers. However, in layer 2/3 we found that the potency of inhibitory synapses from PV-FS INs to PNs and excitatory synapses from PNs to PV-FS INs are enhanced, resulting in greater feedforward inhibition of excitatory input from layer 4. Thus, our data show that *Arid1b* haploinsufficiency results in opposite changes in the ratio of excitation to inhibition (E/I) dependent on cortical layer and cell-type. The reason for this difference is unclear, however, the local niche environments in layers 2/3 and 5 are unique and may differentially guide the development of PV-FS INs (Fishell and Kepecs, 2019). Furthermore, it remains unclear whether these changes are pathological or homeostatic to maintain overall network function (Antoine et al., 2019). Our data provide a framework for future experiments *in vivo* to test specific hypotheses regarding differential changes in E/I balance in *Arid1b* haploinsufficient mice.

### Mechanisms and consequences of disrupted short-term facilitation at excitatory synapses to SST-NFS interneurons

PV-FS INs have been the focus of most research on the inhibitory circuit mechanisms underlying neurodevelopmental disorders. However, pathology in other IN subtypes, such as SST-NFS INs, likely contributes to cognitive and behavioral deficits. SST-NFS INs target the distal dendrites of PNs and play an important role in regulating synaptic integration and network excitability (Kapfer et al., 2007; Silberberg and Markram, 2007). SST-NFS INs receive most of their excitatory synaptic input from neighboring PNs (Naskar et al., 2021). These synapses demonstrate a striking form of short-term plasticity in which EPSCs facilitate in response to trains of presynaptic action potentials (Reyes et al., 1998; Gibson et al., 1999; Losonczy et al., 2002; Sun et al., 2009; Sylwestrak and Ghosh, 2012; Urban-Ciecko et al., 2018; Stachniak et al., 2019). Thus, SST-NFS INs provide feedback inhibition as local excitation increases (Kapfer et al., 2007). We found that *Arid1b* haploinsufficiency results in a novel, cell-autonomous synaptic deficit in which this facilitation is impaired. This is expected to reduce feedback inhibition within cortical circuits to alter network balance and function *in vivo*.

Facilitation of excitatory input is controlled by a postsynaptic mechanism in SST-NFS INs; an axon from a single PN provides a facilitating synapse to an SST-NSF IN and a depressing synapse to a PV-FS IN (Reyes et al., 1998). To generate facilitation, SST-NFS INs express a member of the leucine-rich repeat synaptic proteins called Elfn1 (Sylwestrak and Ghosh, 2012; Tomioka et al., 2014; de Wit and Ghosh, 2016). Expression of Elfn1 postsynaptic to excitatory synapses recruits and binds presynaptic metabotropic glutamate receptor 7 (mGluR7) and activates presynaptic GluK2 subunit-containing kainate receptors (GluK2-KARs) via an unknown mechanism (Sylwestrak and Ghosh, 2012; Tomioka et al., 2014; Stachniak et al., 2019). In cortical layer 2/3, mGluR7 suppresses synaptic vesicle release during initial presynaptic spikes, while activation of GluK2-KARs during subsequent spikes potentiates transmitter release (Sylwestrak and Ghosh, 2012; Stachniak et al., 2019). Together, engagement of presynaptic mGluR7 and GluK2-KARs during PN action potential trains results in strongly facilitating postsynaptic responses in SST-NFS INs (Stachniak et al., 2019). Thus, our data suggest that *Arid1b* haploinsufficiency disrupts Elfn1 expression or function in SST-NFS INs. Interestingly, we found a deficit in synaptic potency but no change in synaptic failure rate. Thus, *Arid1b* haploinsufficiency may selectively disrupt activation of GluK2-KARs. Regardless, Elfn1 may be a novel therapeutic target to correct circuit function in ARID1B-related disorders.

## METHODS

### Animals

All experiments were conducted in accordance with animal protocols approved by the Ohio State University IACUC. Male and female mice were used without bias. For experiments in mature circuits in neocortex, mice were 4 – 10 weeks old. For experiments in neonatal hippocampus, mice were postnatal days 5 – 7. Nkx2.1-Cre, Arid1b(floxed/floxed), and Ai14 mice were obtained from the Jackson Laboratory (stock nos. 008661, 032061, 007914).

### Slice preparation

Mice were anesthetized with isoflurane and then decapitated. The brain was dissected in ice-cold artificial cerebrospinal fluid (ACSF) containing (in mM): 100 sucrose, 80 NaCl, 3.5 KCl, 24 NAHCO3, 1.25 NaH2PO4, 4.5 MgCl, 0.5 CaCl2, and 10 glucose, saturated with 95% O_2_ and 5% CO_2_. Coronal sections of primary visual cortex or horizontal sections of hippocampus (300 μm) were cut using a Leica VT 1200S vibratome (Leica Microsystems) and incubated in the above solution at 35 °C for 30 minutes post-dissection. Slices were then maintained at room temperature until use in recording ACSF containing (in mM): 130 NaCl, 3.5 KCl, 24 NaHCO3, 1.25 NaH2PO4, 1.5 MgCl, 2.5 CaCl2, and 10 glucose, saturated with 95% O_2_ and 5% CO_2_. Mice older than 6 weeks were first anesthetized with intraperitoneal ketamine/xylazine (100mg/kg / 10 mg/kg) and then perfused for 2 minutes with ice-cold sucrose-based ACSF.

### Electrophysiology

For recording, slices were constantly perfused with ACSF at 2 mL/min at a temperature of 31-33 °C. Cells were visualized using an upright microscope (Scientifica SliceScope) with a 40x water-immersion objective (Olympus), camera (Scientifica SciCam Pro), and Oculus software. Retrobeads, rg-AAV-CAG-GFP, and tdTomato-expression were identified by fluorescence (CoolLED pE-300ultra). Recording pipettes were pulled from borosilicate glass (World Precision Instruments) to a resistance of 3-5 MΩ using a vertical pipette puller (Narishige PC-100). Whole-cell patch clamp recordings were acquired using a Multiclamp 700B amplifier (Molecular Devices), filtered at 3 kHz (Bessel filter), and digitized at 10 or 20 kHz (Digidata 1550B and pClamp v11.1, Molecular Devices). Recordings were not corrected for liquid junction potential. Series resistance (10-25 MΩ) was closely monitored, and recordings were excluded from analysis if series resistance passed 25 MΩ. In current-clamp mode, cells were biased to a membrane potential (Vm) of -70 mV; in voltage-clamp mode, a holding potential of -70 mV was applied. The internal solution used to collect intrinsic membrane properties and excitatory postsynaptic currents and potentials contained (in mM): 130 K-gluconate, 5 KCl, 2 NaCl, 4 MgATP, 0.3 NaGTP, 10 phosphocreatine, 10 HEPES, 0.5 EGTA, and 0.2% biocytin (Sigma-Aldrich catalog #B4261). The calculated ECl- for this solution was -79 mV. The internal solution used to record GABA_A_-mediated currents in voltage-clamp at a holding potential of -70 mV contained (in mM): 85 K-gluconate, 45 KCl, 2 NaCl, 4 MgATP, 0.3 NaGTP, 10 phosphocreatine, 10 HEPES, and 0.5 EGTA, for a calculated ECl- of -29 mV. The pH of these two internal solutions was adjusted to 7.4 with KOH. The internal solution for to record inward and outward currents evoked by layer 4 stimulation contained (in mM): 135 CsMeSO4, 8 KCl, 4 MgATP, 0.3 Na2GTP, 5 QX–314, 0.5 EGTA, and 0.2% biocytin. The pH of this internal solution was adjusted to 7.4 with CsOH.

For paired recordings of synaptic connections, presynaptic cells were made to fire trains of action potentials (25 at 50 Hz) using 2 nA steps of 2 ms duration every 10 s for 10 trials. Post-synaptic cells were recorded in voltage-clamp at -70 mV. To analyze postsynaptic currents in paired recordings, we first zeroed the data by subtracting the baseline and then performed 10 repetitions of binomial (Gaussian) smoothing. All postsynaptic currents (PSCs) were then analyzed relative to the timing of the peak of each presynaptic spike during the train. PSCs were detected by threshold crossing (-7 to -10 pA); if this threshold was already crossed at the instant of spike peak, the PSC data associated with that spike of that trial were discarded as being contaminated by spontaneous events. The proportion of failures was calculated as the ratio of evoked PSCs to the total number of non-contaminated trials for each presynaptic spike. The PSC potency was calculated as the average peak amplitude of all successfully evoked PSCs for each presynaptic spike (i.e., failures were not included in the average).

Input resistance (Rin) was measured using a linear regression of voltage deflections (±20 mV from -70 mV resting membrane potential) in response to 1 s current steps. To calculate voltage sag, the membrane potential was biased to -70 mV (V_initial) followed by injection of a 1 s negative current step of sufficient amplitude to reach a steady state Vm of -90 mV during the last 200 ms of the current injection (V_sag). The peak hyperpolarized Vm prior to sag (V_hyp) was used to calculate the sag index as (V_hyp – V_sag)/(V_hyp – V_initial). Firing rate and adaptation ratio were collected from sweeps at twice threshold necessary to evoke an action potential. Adaptation ratio was calculated as the inter-spike interval between the first two action potentials divided by the averaged inter-spike interval between the last four action potentials.

Extracellular stimulation of layer 4 was performed with a constant current isolator (World Precision Instruments A365 Stimulus Isolator) to deliver current through a low resistance (∼750 kΩ) ACSF-filled glass pipette. Pulses were 0.1 ms long and given every 20 seconds, with three pulses per stimulus intensity.

### Stereotaxic injections

For injections of retrograde tracer into the superior colliculus, mice were anesthetized with 5% isoflurane and mounted in a stereotax (Neurostar, Germany). 24 hours prior to surgery, ibuprofen was added to the cage drinking water and maintained for 72 hours post-operation. Prior to surgery, mice were provided with extended release (48 – 72 hour) buprenorphine (1 mg/kg via subcutaneous injection) for additional post-operative analgesia. Green retrobeads (Lumafluor) or retrograde serotype AAV-CAG-GFP (gift from Edward Boyden; Addgene viral prep # 37825-AAVrg) were injected via a glass capillary nanoinjector (Neurostar, Germany) back filled with light mineral oil. The superior colliculus was targeted using the following coordinates: 4.0 (±0.2) mm caudal and 0.4 mm lateral to bregma, and 1.6 mm deep from the dura. 100 nl of retrobeads or 400 nl of AAV were injected at 75 nl/min. The pipette was left in place for 10 min following the injection before removal. For retrobead injections, mice were sacrificed after at least 2 days post-injection; for AAV injections, mice were sacrificed after at least 7 days post-injection.

### Data analysis and statistics

Electrophysiology data were analyzed in Igor Pro v8.04 (WaveMetrics) using custom routines; pClamp files were imported into Igor using NeuroMatic (Rothman and Silver, 2018). Statistical tests were performed using Graphpad Prism v10.1.1 (Graphpad Software, San Diego, CA). Distributions were first tested for normality using a Shapiro-Wilk test. Distributions that did not violate normality were compared using parametric tests (unpaired t test, one-way ANOVA, or two-way ANOVA as appropriate, described in figure legends), while those that did were compared using nonparametric tests (Mann-Whitney U test or Kruskal-Wallis test as appropriate, described in figure legends). Tests of interneuron subtype distributions were performed with a chi-squared test. Post hoc analyses were performed as appropriate and described in figure legends. Data in the text and graphs are reported as mean ± SEM.

## Acknowledgements

This work was supported by funding from the Simons Foundation Autism Research Initiative (SFARI Pilot Award #724187) and NIH (R01MH124870) to J.C.W.

## REFERENCES

Allene C, Cattani A, Ackman JB, Bonifazi P, Aniksztejn L, Ben-Ari Y, Cossart R (2008) Sequential generation of two distinct synapse-driven network patterns in developing neocortex. J Neurosci 28:12851–12863.

Anderson SA, Marin O, Horn C, Jennings K, Rubenstein JL (2001) Distinct cortical migrations from the medial and lateral ganglionic eminences. Development (Cambridge, England) 128:353–363.

Antoine MW, Langberg T, Schnepel P, Feldman DE (2019) Increased Excitation-Inhibition Ratio Stabilizes Synapse and Circuit Excitability in Four Autism Mouse Models. Neuron 101:648–661 e644.

Ariza J, Rogers H, Hashemi E, Noctor SC, Martinez-Cerdeno V (2018) The Number of Chandelier and Basket Cells Are Differentially Decreased in Prefrontal Cortex in Autism. Cereb Cortex 28:411–420.

Bandler RC, Mayer C, Fishell G (2016) Cortical interneuron specification: the juncture of genes, time and geometry. Curr Opin Neurobiol 42:17–24.

Ben-Ari Y (2001) Developing networks play a similar melody. Trends Neurosci 24:353–360.

Ben-Ari Y, Cherubini E, Corradetti R, Gaiarsa JL (1989) Giant synaptic potentials in immature rat CA3 hippocampal neurones. J Physiol 416:303–325.

Ben-Ari Y, Gaiarsa JL, Tyzio R, Khazipov R (2007) GABA: a pioneer transmitter that excites immature neurons and generates primitive oscillations. Physiol Rev 87:1215-1284.

Blankenship AG, Feller MB (2010) Mechanisms underlying spontaneous patterned activity in developing neural circuits. Nat Rev Neurosci 11:18–29.

Brown KN, Chen S, Han Z, Lu CH, Tan X, Zhang XJ, Ding L, Lopez-Cruz A, Saur D, Anderson SA, Huang K, Shi SH (2011) Clonal production and organization of inhibitory interneurons in the neocortex. Science 334:480–486.

Butt SJ, Fuccillo M, Nery S, Noctor S, Kriegstein A, Corbin JG, Fishell G (2005) The temporal and spatial origins of cortical interneurons predict their physiological subtype. Neuron 48:591–604.

Celen C et al. (2017) Arid1b haploinsufficient mice reveal neuropsychiatric phenotypes and reversible causes of growth impairment. Elife 6.

Ciceri G, Dehorter N, Sols I, Huang ZJ, Maravall M, Marin O (2013) Lineage-specific laminar organization of cortical GABAergic interneurons. Nat Neurosci.

Contractor A, Ethell IM, Portera-Cailliau C (2021) Cortical interneurons in autism. Nat Neurosci 24:1648–1659.

de Wit J, Ghosh A (2016) Specification of synaptic connectivity by cell surface interactions. Nat Rev Neurosci 17:22–35.

Denaxa M, Neves G, Rabinowitz A, Kemlo S, Liodis P, Burrone J, Pachnis V (2018) Modulation of Apoptosis Controls Inhibitory Interneuron Number in the Cortex. Cell Rep 22:1710–1721.

Duan ZRS, Che A, Chu P, Modol L, Bollmann Y, Babij R, Fetcho RN, Otsuka T, Fuccillo MV, Liston C, Pisapia DJ, Cossart R, De Marco Garcia NV (2020) GABAergic Restriction of Network Dynamics Regulates Interneuron Survival in the Developing Cortex. Neuron 105:75–92.e75.

Ellegood J, Petkova SP, Kinman A, Qiu LR, Adhikari A, Wade AA, Fernandes D, Lindenmaier Z, Creighton A, Nutter LMJ, Nord AS, Silverman JL, Lerch JP (2021) Neuroanatomy and behavior in mice with a haploinsufficiency of AT-rich interactive domain 1B (ARID1B) throughout development. Mol Autism 12:25.

Filice F, Vorckel KJ, Sungur AO, Wohr M, Schwaller B (2016) Reduction in parvalbumin expression not loss of the parvalbumin-expressing GABA interneuron subpopulation in genetic parvalbumin and shank mouse models of autism. Mol Brain 9:10.

Filice F, Janickova L, Henzi T, Bilella A, Schwaller B (2020) The Parvalbumin Hypothesis of Autism Spectrum Disorder. Front Cell Neurosci 14:577525.

Fishell G, Kepecs A (2019) Interneuron Types as Attractors and Controllers. Annu Rev Neurosci.

Gibson JR, Beierlein M, Connors BW (1999) Two networks of electrically coupled inhibitory neurons in neocortex. Nature 402:75–79.

Godavarthi SK, Sharma A, Jana NR (2014) Reversal of reduced parvalbumin neurons in hippocampus and amygdala of Angelman syndrome model mice by chronic treatment of fluoxetine. J Neurochem 130:444–454.

Goel A, Cantu DA, Guilfoyle J, Chaudhari GR, Newadkar A, Todisco B, de Alba D, Kourdougli N, Schmitt LM, Pedapati E, Erickson CA, Portera-Cailliau C (2018) Impaired perceptual learning in a mouse model of Fragile X syndrome is mediated by parvalbumin neuron dysfunction and is reversible. Nat Neurosci 21:1404–1411.

Harris KD, Shepherd GM (2015) The neocortical circuit: themes and variations. Nat Neurosci 18:170–181.

Harwell CC, Fuentealba LC, Gonzalez-Cerrillo A, Parker PR, Gertz CC, Mazzola E, Garcia MT, Alvarez-Buylla A, Cepko CL, Kriegstein AR (2015) Wide Dispersion and Diversity of Clonally Related Inhibitory Interneurons. Neuron 87:999–1007.

Hashemi E, Ariza J, Rogers H, Noctor SC, Martinez-Cerdeno V (2017) The Number of Parvalbumin-Expressing Interneurons Is Decreased in the Prefrontal Cortex in Autism. Cereb Cortex 27:1931–1943.

Jung EM, Moffat JJ, Liu J, Dravid SM, Gurumurthy CB, Kim WY (2017) Arid1b haploinsufficiency disrupts cortical interneuron development and mouse behavior. Nat Neurosci 20:1694–1707.

Ka M, Chopra DA, Dravid SM, Kim WY (2016) Essential Roles for ARID1B in Dendritic Arborization and Spine Morphology of Developing Pyramidal Neurons. J Neurosci 36:2723–2742.

Kapfer C, Glickfeld LL, Atallah BV, Scanziani M (2007) Supralinear increase of recurrent inhibition during sparse activity in the somatosensory cortex. Nat Neurosci 10:743–753.

Kim H, Kim D, Cho Y, Kim K, Roh JD, Kim Y, Yang E, Kim SS, Ahn S, Kim H, Kang H, Bae Y, Kim E (2022) Early postnatal serotonin modulation prevents adult-stage deficits in Arid1b-deficient mice through synaptic transcriptional reprogramming. Nature communications 13:5051.

Kourdougli N, Suresh A, Liu B, Juarez P, Lin A, Chung DT, Graven Sams A, Gandal MJ, Martínez-Cerdeño V, Buonomano DV, Hall BJ, Mombereau C, Portera-Cailliau C (2023) Improvement of sensory deficits in fragile X mice by increasing cortical interneuron activity after the critical period. Neuron 111:2863–2880.e2866.

Lee AT, Gee SM, Vogt D, Patel T, Rubenstein JL, Sohal VS (2014) Pyramidal neurons in prefrontal cortex receive subtype-specific forms of excitation and inhibition. Neuron 81:61–68.

Losonczy A, Zhang L, Shigemoto R, Somogyi P, Nusser Z (2002) Cell type dependence and variability in the short-term plasticity of EPSCs in identified mouse hippocampal interneurones. J Physiol 542:193–210.

Mayer C, Jaglin XH, Cobbs LV, Bandler RC, Streicher C, Cepko CL, Hippenmeyer S, Fishell G (2015) Clonally Related Forebrain Interneurons Disperse Broadly across Both Functional Areas and Structural Boundaries. Neuron 87:989–998.

Moffat JJ, Smith AL, Jung EM, Ka M, Kim WY (2021a) Neurobiology of ARID1B haploinsufficiency related to neurodevelopmental and psychiatric disorders. Mol Psychiatry.

Moffat JJ, Jung EM, Ka M, Jeon BT, Lee H, Kim WY (2021b) Differential roles of ARID1B in excitatory and inhibitory neural progenitors in the developing cortex. Sci Rep 11:3856.

Moffat JJ, Jung EM, Ka M, Smith AL, Jeon BT, Santen GWE, Kim WY (2019) The role of ARID1B, a BAF chromatin remodeling complex subunit, in neural development and behavior. Prog Neuropsychopharmacol Biol Psychiatry 89:30–38.

Naskar S, Qi J, Pereira F, Gerfen CR, Lee S (2021) Cell-type-specific recruitment of GABAergic interneurons in the primary somatosensory cortex by long-range inputs. Cell Rep 34:108774.

Nomura T (2021) Interneuron Dysfunction and Inhibitory Deficits in Autism and Fragile X Syndrome. Cells 10.

Pagliaroli L, Trizzino M (2021) The Evolutionary Conserved SWI/SNF Subunits ARID1A and ARID1B Are Key Modulators of Pluripotency and Cell-Fate Determination. Front Cell Dev Biol 9:643361.

Pangratz-Fuehrer S, Hestrin S (2011) Synaptogenesis of electrical and GABAergic synapses of fast-spiking inhibitory neurons in the neocortex. J Neurosci 31:10767–10775.

Patz S, Grabert J, Gorba T, Wirth MJ, Wahle P (2004) Parvalbumin expression in visual cortical interneurons depends on neuronal activity and TrkB ligands during an Early period of postnatal development. Cereb Cortex 14:342–351.

Petros TJ, Bultje RS, Ross ME, Fishell G, Anderson SA (2015) Apical versus Basal Neurogenesis Directs Cortical Interneuron Subclass Fate. Cell Rep 13:1090–1095.

Priya R, Paredes MF, Karayannis T, Yusuf N, Liu X, Jaglin X, Graef I, Alvarez-Buylla A, Fishell G (2018) Activity Regulates Cell Death within Cortical Interneurons through a Calcineurin-Dependent Mechanism. Cell Rep 22:1695–1709.

Reyes A, Lujan R, Rozov A, Burnashev N, Somogyi P, Sakmann B (1998) Target-cell-specific facilitation and depression in neocortical circuits. Nat Neurosci 1:279–285.

Rothman JS, Silver RA (2018) NeuroMatic: An Integrated Open-Source Software Toolkit for Acquisition, Analysis and Simulation of Electrophysiological Data. Front Neuroinform 12:14.

Santen GW, Clayton-Smith J (2014) The ARID1B phenotype: what we have learned so far. Am J Med Genet C Semin Med Genet 166C:276-289.

Silberberg G, Markram H (2007) Disynaptic inhibition between neocortical pyramidal cells mediated by Martinotti cells. Neuron 53:735–746.

Smith AL, Jung EM, Jeon BT, Kim WY (2020) Arid1b haploinsufficiency in parvalbumin- or somatostatin-expressing interneurons leads to distinct ASD-like and ID-like behavior. Sci Rep 10:7834.

Sohal VS, Rubenstein JLR (2019) Excitation-inhibition balance as a framework for investigating mechanisms in neuropsychiatric disorders. Mol Psychiatry 24:1248–1257.

Southwell DG, Paredes MF, Galvao RP, Jones DL, Froemke RC, Sebe JY, Alfaro-Cervello C, Tang Y, Garcia-Verdugo JM, Rubenstein JL, Baraban SC, Alvarez-Buylla A (2012) Intrinsically determined cell death of developing cortical interneurons. Nature 491:109–113.

Spitzer NC (2006) Electrical activity in early neuronal development. Nature 444:707–712.

Stachniak TJ, Sylwestrak EL, Scheiffele P, Hall BJ, Ghosh A (2019) Elfn1-Induced Constitutive Activation of mGluR7 Determines Frequency-Dependent Recruitment of Somatostatin Interneurons. J Neurosci 39:4461–4474.

Stevens CF, Wang Y (1995) Facilitation and depression at single central synapses. Neuron 14:795–802.

Sun HY, Bartley AF, Dobrunz LE (2009) Calcium-permeable presynaptic kainate receptors involved in excitatory short-term facilitation onto somatostatin interneurons during natural stimulus patterns. J Neurophysiol 101:1043–1055.

Sylwestrak EL, Ghosh A (2012) Elfn1 regulates target-specific release probability at CA1-interneuron synapses. Science 338:536–540.

Tang X, Jaenisch R, Sur M (2021) The role of GABAergic signalling in neurodevelopmental disorders. Nat Rev Neurosci 22:290–307.

Tasic B et al. (2018) Shared and distinct transcriptomic cell types across neocortical areas. Nature 563:72–78.

Tomioka NH, Yasuda H, Miyamoto H, Hatayama M, Morimura N, Matsumoto Y, Suzuki T, Odagawa M, Odaka YS, Iwayama Y, Won Um J, Ko J, Inoue Y, Kaneko S, Hirose S, Yamada K, Yoshikawa T, Yamakawa K, Aruga J (2014) Elfn1 recruits presynaptic mGluR7 in trans and its loss results in seizures. Nature communications 5:4501.

Tremblay R, Lee S, Rudy B (2016) GABAergic Interneurons in the Neocortex: From Cellular Properties to Circuits. Neuron 91:260–292.

Urban-Ciecko J, Jouhanneau JS, Myal SE, Poulet JFA, Barth AL (2018) Precisely Timed Nicotinic Activation Drives SST Inhibition in Neocortical Circuits. Neuron 97:611–625 e615.

van der Sluijs PJ et al. (2019) The ARID1B spectrum in 143 patients: from nonsyndromic intellectual disability to Coffin-Siris syndrome. Genet Med 21:1295–1307.

Wang W, Côté J, Xue Y, Zhou S, Khavari PA, Biggar SR, Muchardt C, Kalpana GV, Goff SP, Yaniv M, Workman JL, Crabtree GR (1996) Purification and biochemical heterogeneity of the mammalian SWI-SNF complex. The EMBO journal 15:5370–5382.

Wang X, Nagl NG, Wilsker D, Van Scoy M, Pacchione S, Yaciuk P, Dallas PB, Moran E (2004) Two related ARID family proteins are alternative subunits of human SWI/SNF complexes. Biochem J 383:319–325.

Wester JC, McBain CJ (2016) Interneurons Differentially Contribute to Spontaneous Network Activity in the Developing Hippocampus Dependent on Their Embryonic Lineage. J Neurosci 36:2646–2662.

Wichterle H, Turnbull DH, Nery S, Fishell G, Alvarez-Buylla A (2001) In utero fate mapping reveals distinct migratory pathways and fates of neurons born in the mammalian basal forebrain. Development (Cambridge, England) 128:3759–3771.

Wong FK, Bercsenyi K, Sreenivasan V, Portales A, Fernandez-Otero M, Marin O (2018) Pyramidal cell regulation of interneuron survival sculpts cortical networks. Nature 557:668–673.

Xu Q, Tam M, Anderson SA (2008) Fate mapping Nkx2.1-lineage cells in the mouse telencephalon. J Comp Neurol 506:16–29.

